# Long-term p21 and p53 trends regulate the frequency of mitosis events and cell cycle arrest

**DOI:** 10.1101/2021.08.17.456721

**Authors:** Anh Phong Tran, Christopher J. Tralie, Caroline Moosmüller, Zehor Belkhatir, José Reyes, Arnold J. Levine, Joseph O. Deasy, Allen R. Tannenbaum

## Abstract

Radiation exposure of healthy cells can halt cell cycle temporarily or permanently. In this work, two single cell datasets that monitored the time evolution of p21 and p53, one subjected to gamma irradiation and the other to x-ray irradiation, are analyzed to uncover the dynamics of this process. New insights into the biological mechanisms were found by decomposing the p53 and p21 signals into transient and oscillatory components. Through the use of dynamic time warping on the oscillatory components of the two signals, we found that p21 signaling lags behind its lead signal, p53, by about 3.5 hours with oscillation periods of around 6 hours. Additionally, through various quantification methods, we showed how p21 levels, and to a lesser extent p53 levels, dictate whether the cells are arrested in their cell cycle and how fast these cells divide depending on their long-term trend in these signals.

## 2 Introduction

The tumor suppressor p53 is often referred to as the “guardian of the genome” for its importance in regulating a large number of target genes. It has an essential role in regulating the cell cycle, inducing cell apoptosis or DNA repair, and senescence [18]. Because of the centrality of p53 in maintaining the integrity of the DNA and preventing mutated cells from proliferating, mutation to the *TP53* gene is found in over 50% of cancers [28]. Inactivation of p53 can take various forms: viral infection, deletion of the p14^*ARF*^ gene, deletion of the carboxy-terminal domain, multiplication of the murine double minute 2 gene (Mdm2), and amino-acid-changing mutation in the DNA-binding domain [34].

The interaction between p53 and its negative regulator Mdm2 is of great importance in understanding its dynamic behavior. The activation of p53 as a result of cell stress or damage induces an increase in Mdm2 levels that in terms inactivates p53 through proteasomal-dependent degradation [16]. Because this feedback loop has a time delay of 30-40 min between the increase in p53 and the increase in Mdm2 transcription, this creates the oscillatory behavior that is often observed in the p53 dynamics [17]. This p53–Mdm2 interaction and the oscillatory dynamics of p53 have been the subject of extensive research [1, 35, 38, 14, 3, 27]. Investigating the p53 pathway and how it relates to cell fate, such as senescence and apoptosis, is essential to developing cancer treatments [31]. In addition to Mdm2, the interactions of p53 with other elements of the pathway such as p14^*ARF*^, Chk2, Wip1, and ATM are considered of importance [3, 9, 13].

Cellular senescence as well as temporary instances of cell cycle arrest is also of great interest as a potential therapeutic target through the prevention of cancerous cells from proliferating [37, 11, 20]. Notably, for cells to remain arrested, this signaling pathway needs to be maintained [4, 12, 23]. Among a large number of targets that p53 influences, the cyclin-dependent kinase (CDK) inhibitor p21 plays a crucial role in the ability of cells to undergo cell cycle arrest [15, 36, 7]. p21 interacts with CDK2 as a bistable switch with increased levels of p21 leading to cell cycle arrest, while low levels of p21 allow cells to escape G1 phase arrest [6, 21, 2, 26, 23].

Through the applications of various mathematical techniques to two single cell datasets that monitored p21 and p53 activity after exposure to various levels of gamma or x-ray radiation, the influence of p21 and p53 signaling on cell mitosis is analyzed.

## 3 Methods

### 3.1 Clustering of periodic signals using t-SNE and the Wasserstein distance

Various types of distance metrics are used when clustering data using t-Distributed Stochastic Neighbor Embedding (t-SNE) [30]. Common implementations include the application of the Euclidean distance, Chebychev distance, correlation, cosine, and Spearman’s rank correlation. By computing pairwise distances between high-dimensional points in the data, the algorithm utilizes this information to compute 2D or 3D representations of the data in a way that aims to cluster the data in a meaningful, interpretable way. Thus, choosing the appropriate distance to employ in the t-SNE framework is of great importance for the quality of the clustering that one derives for the separation of the various subsets of a given dataset.

For time series data, each dimension represents the value of the measured variable at a given time *t*. In the case of oscillating signals such as p21 and p53, employing t-SNE with a distance that compares time series by considering differences across measurements taken at the same time *t* may not yield meaningful clusters. This is because periodic signals may not be synchronized in their phases. Thus, even if one was to compare almost identical cells, slight phase offsets may be enough to make this comparison challenging. This is not as much of a problem with p21 as it is for p53, since the trend in the data is much larger in magnitude than the oscillatory part.

To overcome the limitation in dealing with oscillatory signals, we use the Wasserstein distance (WD) from optimal mass transport (OMT). Through the WD, one considers time series as distributions, removing the dependence on time *t*. Thanks to its properties and natural ability to capture geometric information when comparing different types of data and also the development in computational tools, the WD has attracted widespread interest in many application fields during the last several years. We will just sketch the relevant definitions here. For full details and numerous references, we refer the interested reader to [22, 32, 33].

The ***Wasserstein-1 distance*** (Kantorovich formulation) between two *d*-dimensional probability distributions

*ρ* _0_, *ρ*_1_ defined on a subdomain Ω *⊂* ℝ^*d*^ is given as follows:

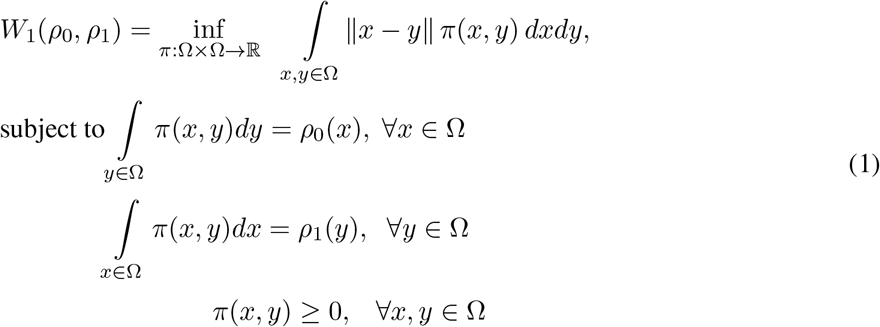

where ║·“ ║is the ground metric. Thus the infimum is taken over all joint distributions *π* : Ω × Ω →ℝ whose marginals are *ρ*_0_, *ρ*_1_.

For the time series analysis of p21 and p53 in the present work, one has *d* = 1, and in this case, the Wasserstein-1 distance has the following very simple form:

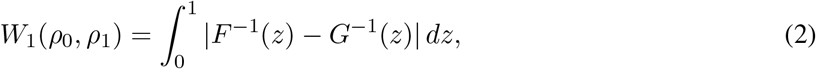

where *F* and *G* denote the cumulative distribution functions of the distributions *ρ*_0_ and *ρ*_1_, respectively.

### 3.2 Analyzing delay between oscillatory signals

#### Signal detrending and amplitude normalization

The preprocessing step for analyzing the relationship between p53 and p21 signals was to detrend the data. This can be done by fitting polynomials, preferably of low order, to the time series data. By finding an appropriate order that captures the trend of the data, one can then subtract the “trend” from the actual data to obtain the oscillatory component. In dealing with signals like p53 and p21 that can be subjected to abrupt changes when the cells are about or are undergoing division, the detrending should be applied to subsets of the overall time series data that avoids including these abrupt changes.

We were able to obtain cleaner results using a method introduced in our previous work [19]. The first part of this technique called the ***Detrended Autocorrelation Periodicity Scoring*** (DAPS) algorithm focuses on the detrending and amplitude normalization using a sliding window embedding, which can be briefly summarized in the following steps:

1. Given the sliding window of length *M* of an *N* -length time series *x*, the sliding windows are arranged into the columns of an *M ×* (*N − M* + 1) matrix *X*, so that the *i*^th^ column of *X* is *SW*_*M*_ [*x*]_*i*_.

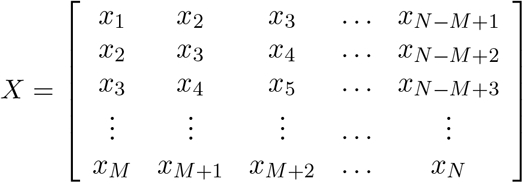 In other words, the *i*^*th*^ skew-diagonal of *X* is a constant value equal to the *i*^th^ value of the time series *xi*. In this work, *M* is chosen to be 11 data points, or 5.5 hours, which is roughly the length of one period of p53 [14].
2. The mean of each column of *X* is subtracted from each element of that column. This normalizes for linear drift in the time series.
3. After point-centering, each column is normalized to be of unit norm. This controls for changes in amplitude in the signal.
4. Lastly, the following operation is performed on the matrix

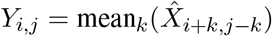

with

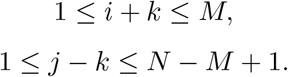

Finally, the *i*^th^ value of the time series *y* is then simply the value of any of the elements in the *i*^th^ skew diagonal of *Y* .

#### Quantifying delay using dynamic time warping and cross-correlation

Dynamic time warping (DTW) [25] is a method that measures the similarity between two time series by finding an optimal time-ordered correspondence, known as a “warping path,” that best maps every index in the first time series to every index in the second time series [5]. Once the signals have been properly detrended and normalized, we apply DTW to determine which of the proteins p53 or p21 leads or lags, as well as the actual delay between the two sequences. It is likely that, because of the periodic nature of these signals, the algorithm may pick up the real delay as well as this value adjusted by integer multiples of the period of these oscillations. We discuss this point in more detail in the **Results** section. The “warping path” of a pure delay is expected to be a straight line in this analysis with the true delay being a stronger signal than the “secondary” alignments that are offset by a certain amount of periods.

Similarly, cross-correlation may be used to determine a correspondence between two time series by shifting one relative to the other. This work introduces the concept of using DTW as an alternative to cross-correlation to capture signal delays, with the hope that other applications may benefit from it even though similar results were obtained using both techniques.

### 3.3 Long-term trend in the signals and occupancy density

The analysis of long-term trends in the signal is performed by using moving averages (m.a.) [8]. By removing the short-term fluctuations, this transformation acts as a low-pass filter that allows highlighting the trend from the overall signal. Using a relatively small window of 9 data points or 4.5 hours, all the time series shown in this work could be cleanly smoothed out. Since the amplitude of the oscillations can be quite large relative to the long-term trend, especially for p53, previous studies focused greatly on describing the behavior of the oscillations. A primary argument of this work is to show that focusing on the trend and not the oscillations for p21 and p53 provide important and novel insights about the current state of a cell as it pertains to cell cycle arrest.

Using moving average values of p21 and p53, 2-D maps of occupancy density were constructed to better understand where these values tend to lie with respect to their likelihood of undergoing mitosis or treatment conditions. By constructing a vector of these moving averages for all time points, each time point then takes up a (*x, y*) coordinate in space based on (*p*53 *m.a., p*21 *m.a.*). In this work, a grid was delimited by approximately the minimum and maximum values for each of the axis. Due to the large discrepancy of low and high values as well as the exponential nature of chemical reactions, a *log*_10_ (*p*21 *m.a.*) vs. *log*_10_ (*p*53 *m.a.*) was used and discretized by 0.1 increment on this scale. This discretized grid is used to count each time a coordinate lies within one of the cells. This is then normalized into the final occupancy density maps.

## 4 Results

### 4.1 Experimental single cell datasets and general trends

The analysis in this work is based on two datasets of individual human cells that were published in [23, 29]. The cells were monitored using fluorescence live-cell imaging for about a week after being subjected to irradiation. The single cells in the first dataset were subjected to gamma irradiation at 0 Gy, 2 Gy, 4 Gy, and 10 Gy. The culture media was replenished daily, except for a subset of the cells subjected to 10 Gy irradiation that had their culture media preserved throughout the experiment. Fluorescence markers for p21 and p53, as well as the mitosis events, were tracked throughout the experiment. The single cells in the second dataset were subjected to x-ray irradiation at 0 Gy, 0.5 Gy, 1 Gy, 2 Gy, 4 Gy, and 8 Gy. Fluorescence markers for p21, p53, and geminin, as well as mitosis events, were monitored throughout the experiment. The values used in this work are fluorescence intensities (a.u.) of p53-mNeonGreen reporter for p53, the mKate2 protein to form a p21-mKate2 complex, and the CFP-hGeminin(1-110) reporter for cell cycle progression [24].

The time series data is presented in Fig. 1 for the single cells exposed to various grades of gamma radiation and in Fig. 2 for cells exposed to x-ray radiation. As shown in Fig. 1(a) and Fig. 2(a), between two-thirds and three-quarters of the unirradiated cells divide 3-5 times during the observation period of 5 days. As the radiation levels increase from 0 to 0.5 to 1 Gy in the x-ray dataset, the fraction of cells that divide often progressively diminishes and is mostly replaced by cells that do not divide or divide only once. The division results for gamma radiation levels of 4 Gy and x-ray radiation of 2 Gy are very similar, indicating the possible more harmful nature of x-ray radiation at equivalent exposure levels that translate into cells dividing less. Due to the two datasets being exposed to different radiation sources and the difference in intensity of the fluorescent light source at the time of the experiment, we reasoned that we would analyze these datasets independently and we focused on results that were conserved between the experimental conditions.

**Figure 1:**
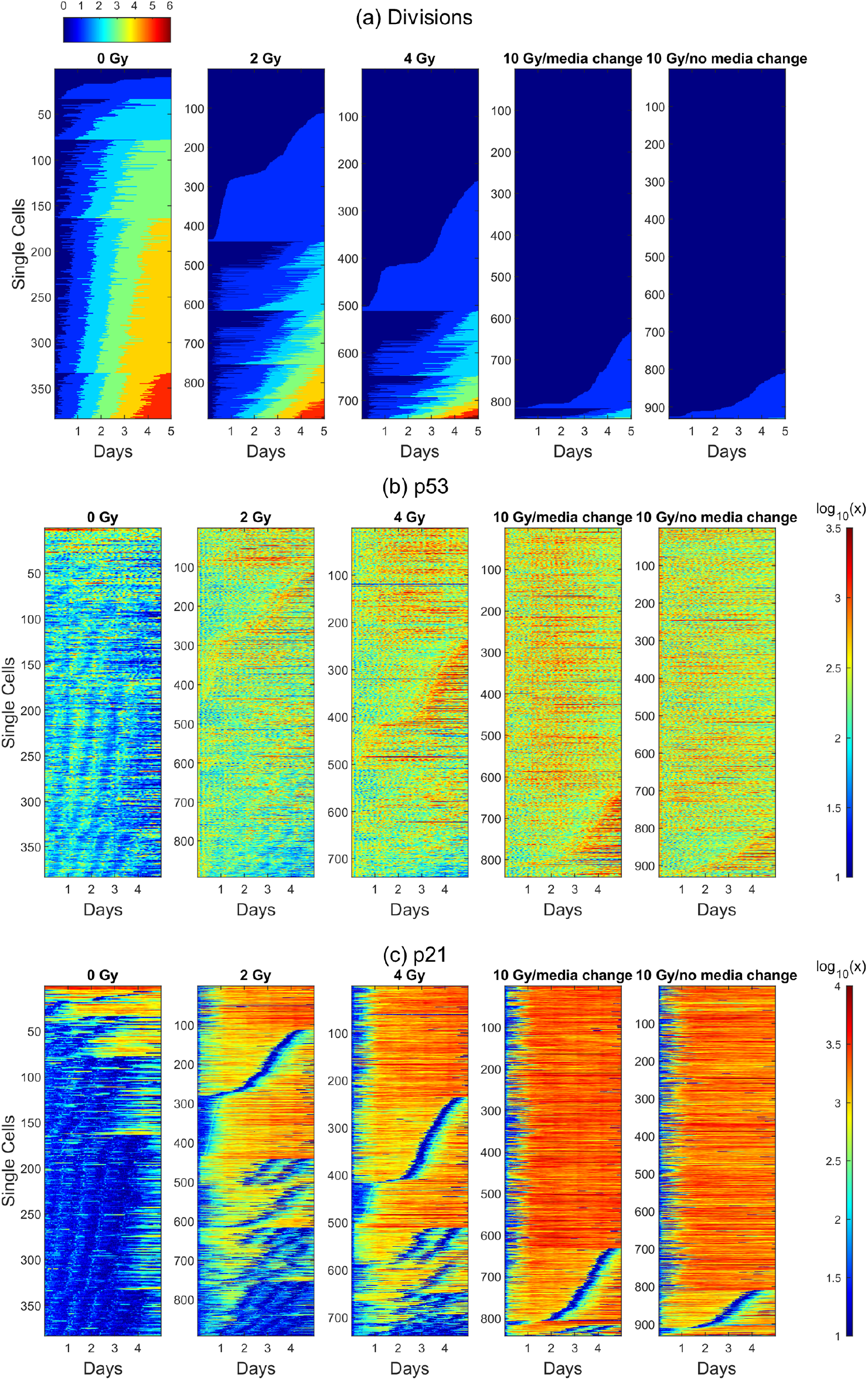
Visualization as a function of time of the gamma irradiation dataset at 0 Gy, 2 Gy, 4 Gy, 10 Gy, and 10 Gy for which the media was removed and replaced by new media. (a) shows the progression of the cells through cell division. The expression levels are then shown on a *log*_10_ scale and broken down by conditions for (b) p53 and (c) p21. The ordering of the single cells is preserved across the different panels for each treatment condition.

**Figure 2:**
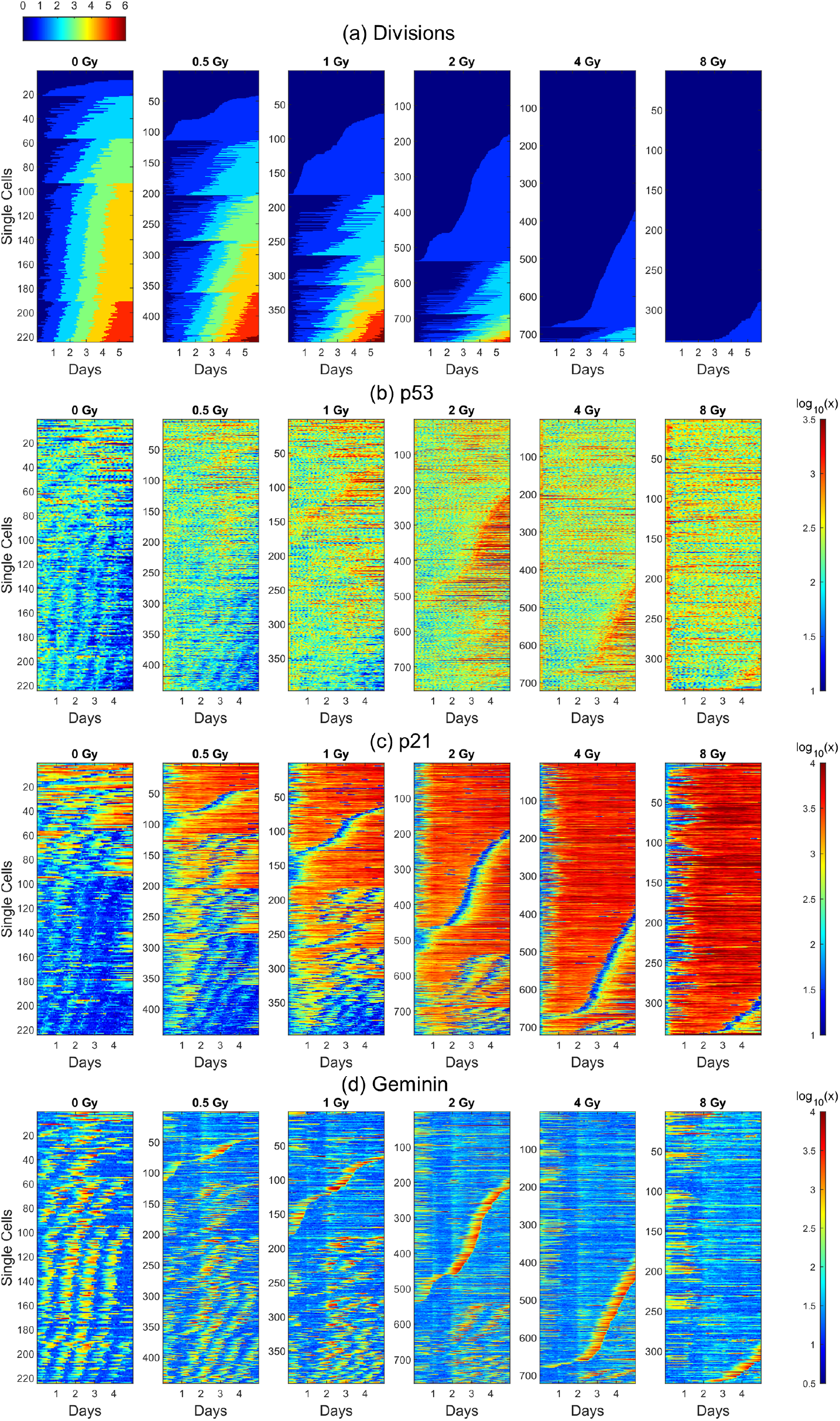
Visualization as a function of time of the x-ray irradiation dataset at 0 Gy, 0.5 Gy, 1 Gy, 2 Gy, 4 Gy, and 8 Gy. (a) shows the progression of the cells through cell division. The expression levels are then shown on a *log*_10_ scale and broken down by conditions for (b) p53, (c) p21, and (d) geminin. The ordering of the single cells is preserved across the different panels for each treatment condition.

Looking at panels Fig. 1(b)-(c) and Fig. 2(c)-(d), p53 levels can be seen to be uniformly oscillating in irradiated cells with the exceptions of cells that divide only once during the 5 days and at least 3 days after radiation exposure. These cells tend to see dramatic increases in p53 levels observed after undergoing their sole mitosis event, consistent with previous reports [29]. On the other hand, p21 levels, exhibit much lower amounts of oscillations, and the trends in the data are of much greater importance to the local fluctuations. From Fig. 2(c) and (d), the relationship is clear that that the upticks of CFP-hGeminin(1-110) correlates with temporal drops in p21 levels. This is because this geminin reporter only accumulates during the S/G_2_/M parts of the cell cycle [30] and p21 is degraded in the G1/S transition [6, 10]. Besides these “gaps” in p21 during the G1/S phases, p21 levels are extremely low in unirradiated cells. p21 levels also rapidly converge to very high values for cells that undergo cell cycle arrest and never divide in the course of the experiment.

### 4.2 p21 and p53 dynamics cluster by number of total observed divisions rather than radiation levels

In the interest of classifying the dynamics of p21 and p53 time series, the t-SNE classification using the 1-Wasserstein distance was applied to the data to generate Fig. 3(a)-(b) for gamma radiation and Fig. 4(a)-(b) for x-ray radiation. The particularity of this distance is that it takes away the temporal aspect of the data and compares exclusively the distributions in p21 or p53 levels between the different time series of individual cells.

**Figure 3:**
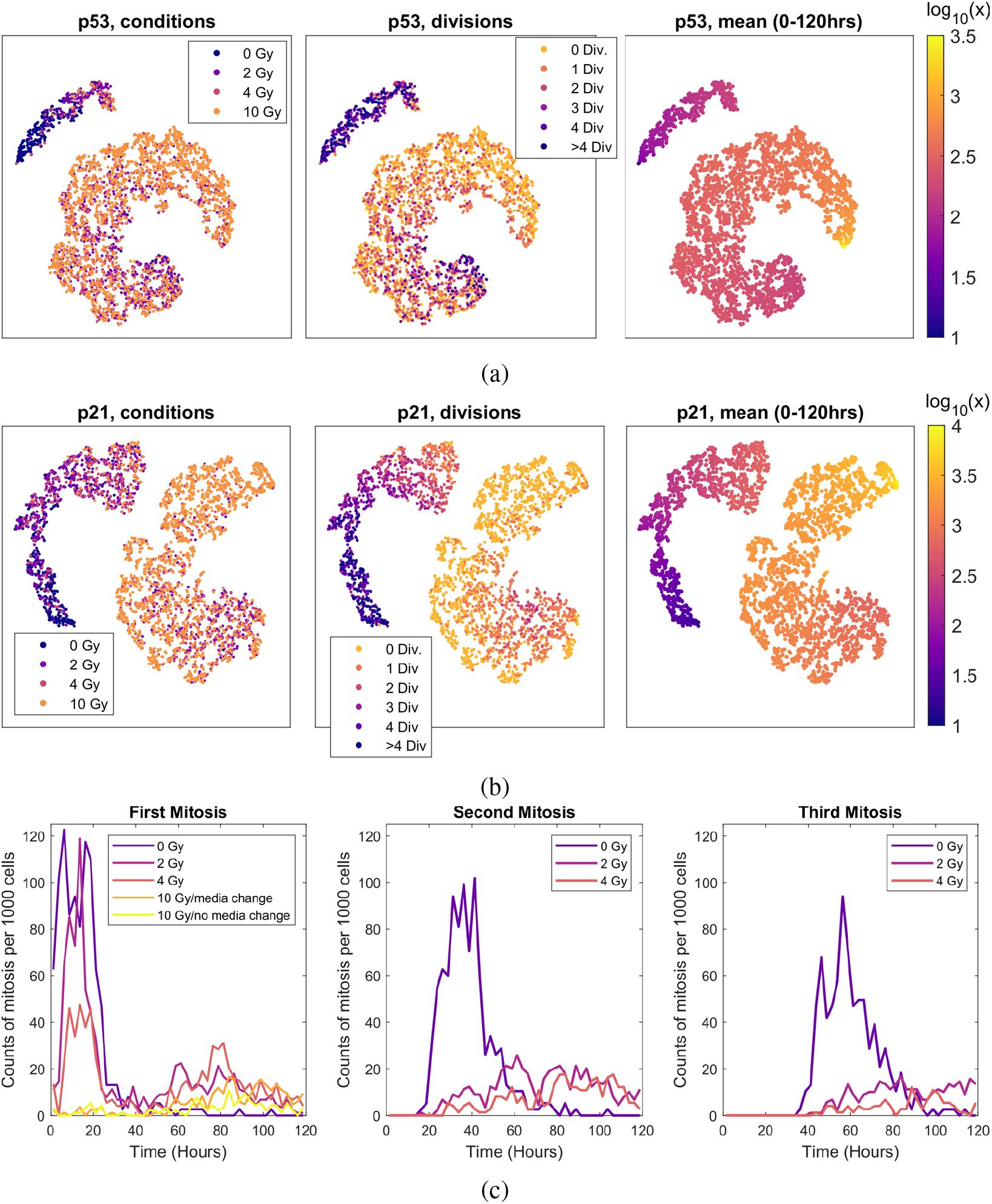
Classification of the gamma irradiation by t-SNE and mitosis events. In (a) and (b), t-SNE based on the Wasserstein-1 distance is applied to the gamma irradiation dataset for the p53 and p21 expression levels. The values of perplexity and exaggeration were picked to be 30 and 4, respectively. For each case, the position of the cells classified by the t-SNE were overlayed by treatment conditions, number of divisions experienced over five days, and the mean values of p21 or p53 during that time. In (c), the divisions are shown as a function by separating first, second, and third mitosis.

**Figure 4:**
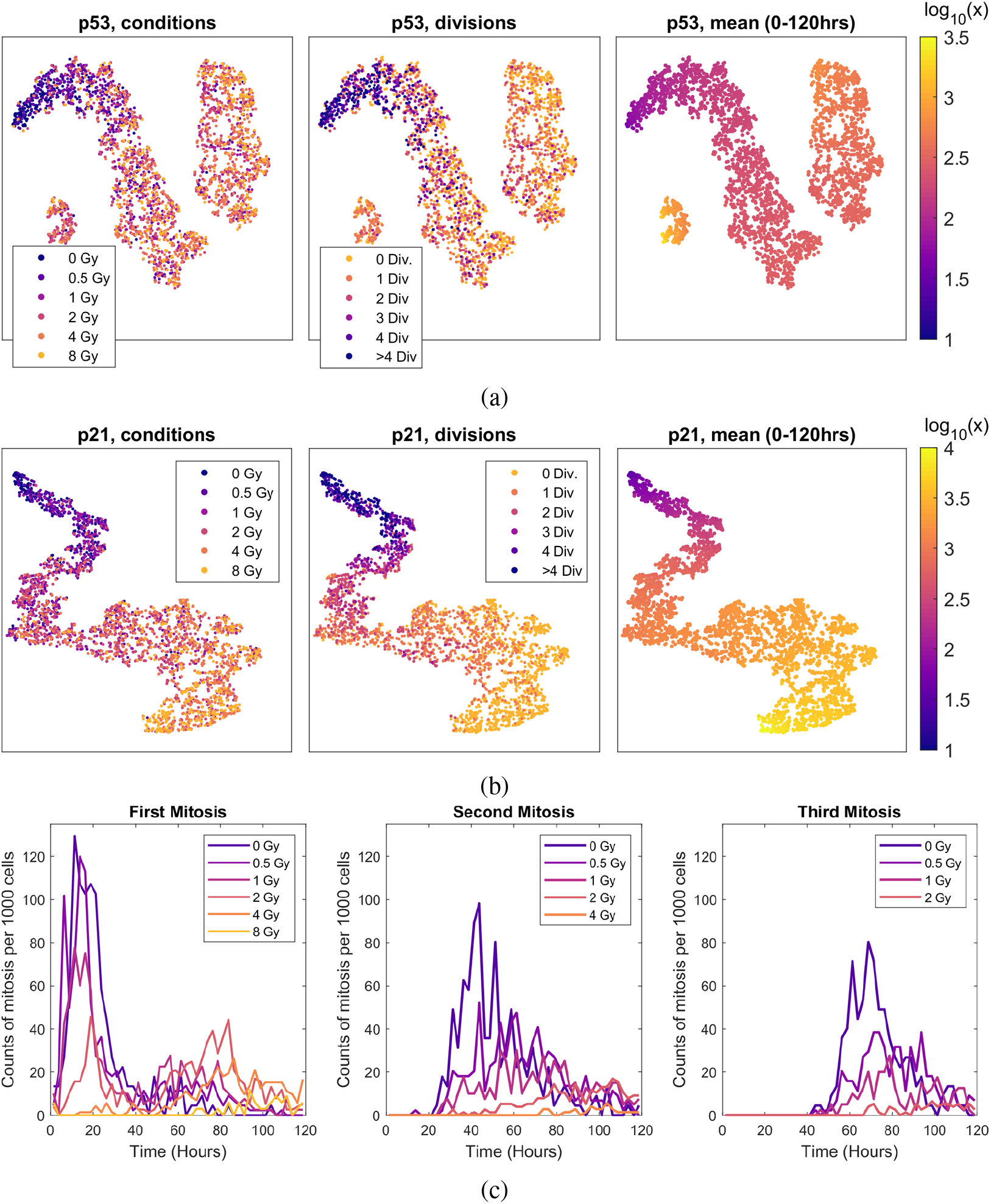
Classification of the x-ray irradiation by t-SNE and mitosis events. In (a) and (b), t-SNE based on the Wasserstein-1 distance is applied to the x-ray irradiation dataset for the p53 and p21 expression levels. The values of perplexity and exaggeration were picked to be 30 and 4, respectively. For each case, the position of the cells classified by the t-SNE were overlayed by treatment conditions, number of divisions experienced over five days, and the mean values of p21 or p53 during that time. In (c), the divisions are shown as a function by separating first, second, and third mitosis.

For each dataset, we matched the t-SNE results on the dynamics with the appropriate labels into four subplots: p53 dynamics by levels of radiation exposure, p53 dynamics by the number of total observed divisions, p21 dynamics by levels of radiation exposure, and p21 dynamics by the number of total observed divisions. From these results, the relation between p21 dynamics and the number of total observed divisions creates the best separation between the groups in the data. The relationship between p21 dynamics and levels of radiation exposure is still significant, but there are a lot of more overlaps between these subcategories of cells. Similarly, the data clusters better relating p53 dynamics to total observed divisions than p53 dynamics to treatment conditions. However, it is clear that considering p53 and p21, the latter is the better indicator as to whether a cell decides to arrest, or divide less often than what one would expect from a healthy cell.

The most meaningful result from the t-SNE classification is that the total number of observed divisions is inversely correlated with the mean values of observed p21 and p53. The correspondence between a lower frequency of observed mitosis events and higher levels of mean p21 levels is particularly striking in the data. This correspondence is not as clear in p53, thus implying that high levels of p53 usually correspond to a lower number of divisions, but that there are also other factors a cell must consider in making the decision to undergo cell cycle arrest. Thus, p21 signaling, being a more downstream indicator, is a cleaner reflection of the cell’s decision to undergo cell cycle arrest.

High levels of irradiation such as 8 Gy for the x-ray dataset and 10 Gy for the gamma-ray dataset do contain mostly cells that have high levels of p21 and p53. These cells tend to either not divide or divide only one time. For cells that are treated with “medium” levels of radiation, a fraction behaves similarly to healthy cells and the rest behaves like damaged cells that were subjected to “high” levels of radiation. Only the healthy cells that were not subjected to radiation damage separate cleanly from the rest of the data. The existence of healthy behaving cells in irradiated conditions is most easily captured in Figs. 3(c) and 4(c). There is almost always a number of cells that divide normally within 20-30 hours of irradiation exposure similar to unirradiated cells at lower levels of radiation. This attests to the heterogeneous nature of the cells when exposed to radiation and the reason why clustering cells by levels of radiation exposure does not cleanly separate the data into clusters. As expected, the ratio of damaged cells compared to healthy cells increase with increased irradiation exposure.

### 4.3 Quantifying the delay between p21 and p53 signaling

Given that p21 lies downstream of the p53 signaling pathway, part of the observed fluctuations in p21 is directly influenced by the oscillatory component of p53. By detrending and normalizing the two signals using part of the DAPS method as shown in Fig. 5(a) for the gamma radiation dataset, these transformed signals were used to calculate the delay between the two signals by warping or “aligning” them to one another using DTW. By performing this step for all time series in the gamma radiation dataset, the aggregated warping paths are shown in Fig. 5(b). There are mostly two main visible straight lines that cut across the graph, one indicating that the p21 signaling is lagging p53 signaling by about 3.5 hours and the other indicating that p21 leads p53 by 2.5 hours. Because the two signals are periodic, the later result is a natural artifact that arises from the fact that the patterns of transformed signals repeat themselves. Thus, the “correct alignment” offset by full periods of the signal will yield local maximum in terms of correlation between the compared signals.

**Figure 5:**
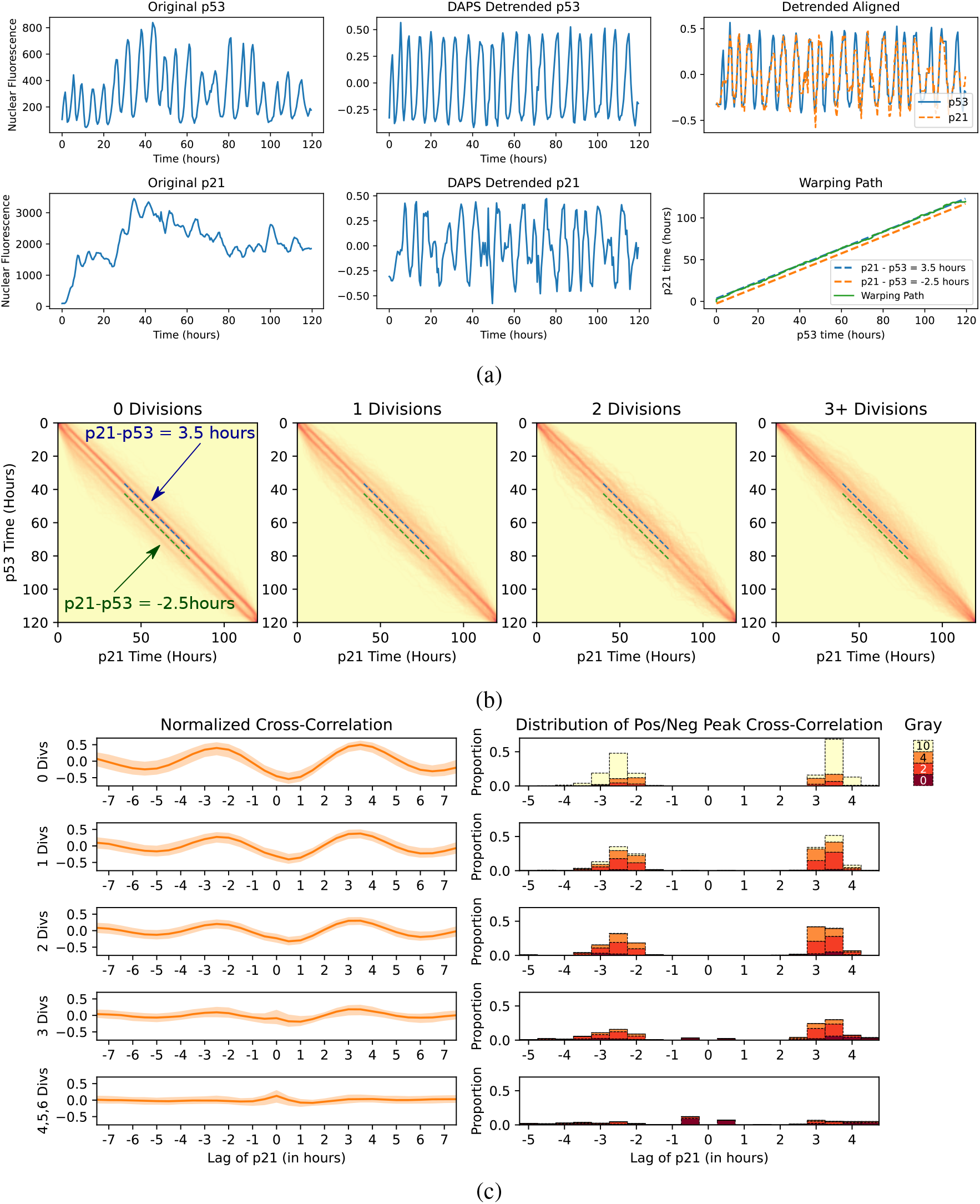
The top figure shows histograms of the parameterizations uncovered via dynamic time warping between the detrended p53 and p21 time series, as a function of number of divisions. The bottom figure shows normalized cross-correlation between the two detrended time series. In both cases, p53 leads p21 by 3.5 hours or lags p21 by 2.5 hours more often for fewer divisions, which is consistent with the period of about 6 hours of p53 during cell damage.

Both the normalized cross-correlation and DTW results in Fig. 5(b)-(c) showed a stronger correlation when aligning with p53 leading p21 by about 3.5 hours. This holds whether looking at cells that did not divide, or divided 1, 2, or 3 times during the 5 days following radiation exposure. This is also identical to trends observed in similar results generated for the x-ray radiation dataset that are shown in Fig. S3. These observations do not hold when looking at cells that divide 3+ times over 5 days whether one looks at the gamma or x-ray radiation datasets. This can be partially explained by p21 signaling being suppressed during the G1/S phases, which represent a major fraction of the cell cycle, reducing the ability to compare the two signals. The oscillations are also noisier when cells are dividing at the pace of healthy cells. Thus, these single cell datasets containing a majority of cells for which the rate of mitosis was slowed down or for which cells are arrested made this analysis on the delay most appropriate.

### 4.4 Long-term p21 and p53 trends play a key role in determining cell behavior vis-à-vis of mitosis

In Fig. 6(a) for the gamma radiation dataset and Fig. S1(a) for the x-ray radiation dataset, using a moving average (m.a.) window of 4.5 hours, p21 m.a. values are plotted as a function of their respective p53 m.a. values at *t* = 0, 12.5, 25, and 50 hours. After 50 hours, the m.a. values of p21 and p53 remain fairly consistent. p53 values settle in a matter of hours as opposed to 1 to 2 days for the p21 values after irradiation. Videos of the datasets that show the evolution at every measured time point over 5 days are available at https://github.com/phongatran/p21p53/. There is a slight long-term drift in the two signals that makes the data points slowly disperse, which may potentially be due to a loss in fluorescence signal over time.

**Figure 6:**
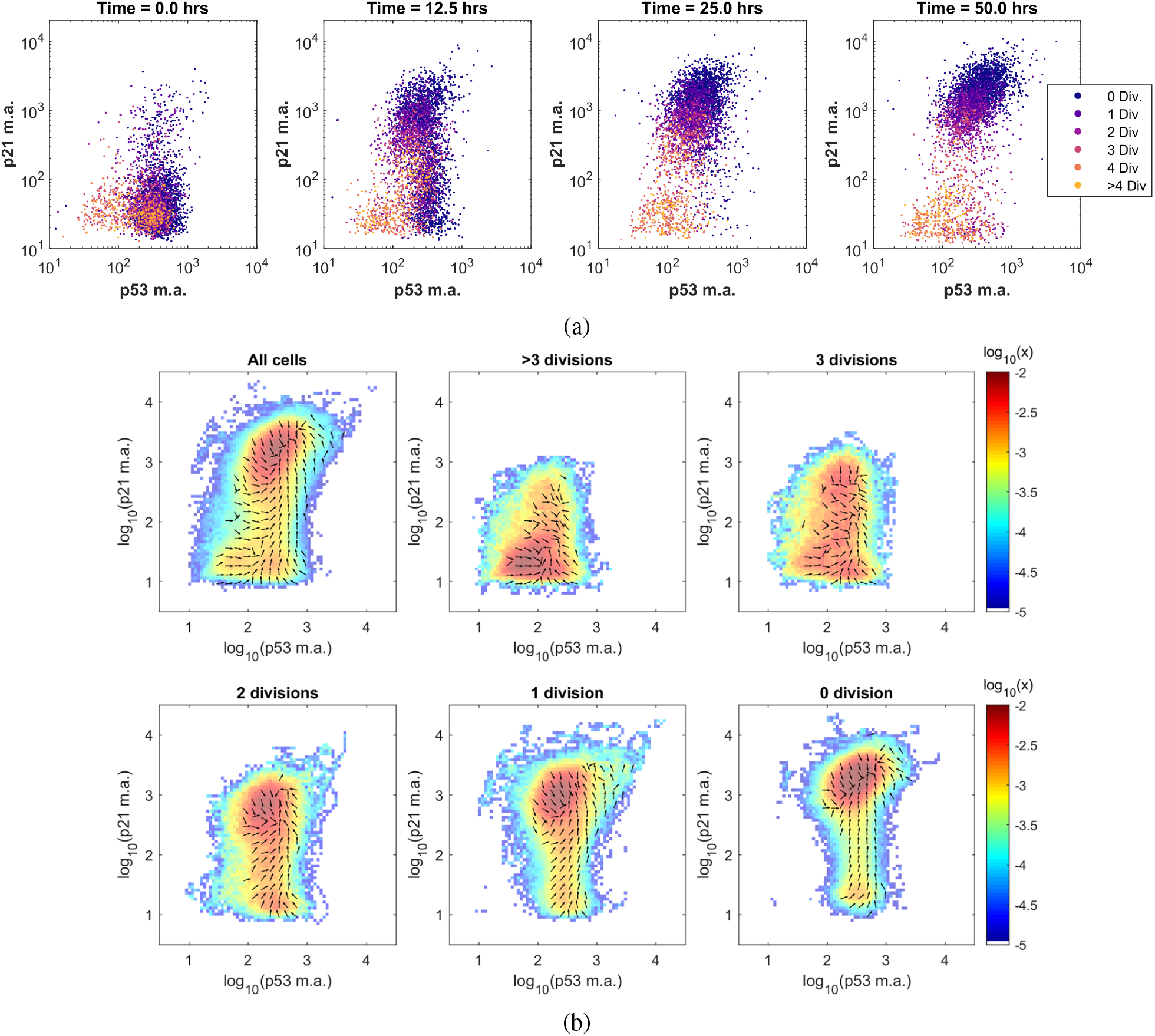
2D representations of the p21 and p53 temporal data for the gamma radiation dataset. (a) Time lapses of the single cells of the moving average values of p21 vs. p53 at different time points of 0, 12.5, 25, and 50 hours after irradiation. The panels are broken down either by number of total divisions over five days or by irradiation conditions. (b) Occupancy density of the cells with a breakdown by the total number of undergone divisions over five days. The numerical value shown in color indicates the probability for a cell to occupy a given square of 0.1 by 0.1 in the log_10_(p21 m.a.) vs. log_10_(p53 m.a.) space in a given hour.

By aggregating the m.a. values of these two markers, the occupancy densities are shown in Fig. 6(b) for the gamma radiation dataset and Fig. S1(b) for the x-ray radiation dataset. From these depictions, it is clear that p21 levels is the main determinant as to how often cells divide over the course of 5 days after irradiation. Note that the cluster of cells moves progressively upwards with the p21 m.a. as the y-axis, looking from *>* 3 down to 0 divisions. Additional plots are available in Fig. S2, illustrating the occupancy densities broken down by radiation levels for both datasets.

To further understand the relationship between the trends in p21 and p53 with respect to cell cycle arrest, the m.a. values are centered around mitosis in Fig. 7 for cells that divide and are exposed to x-ray radiation. In Fig. 7(a), the cells that do not divide have a characteristic jump in p53 m.a. values that then subside within the first few hours after the cell receives the irradiation damage. The p21 m.a. values are constantly increasing for the next 20-30 hours until they reach a plateau of about ∼3000-5500. These non-dividing cells remain in this state with little variation in geminin, p21, or p53 m.a. values.

**Figure 7:**
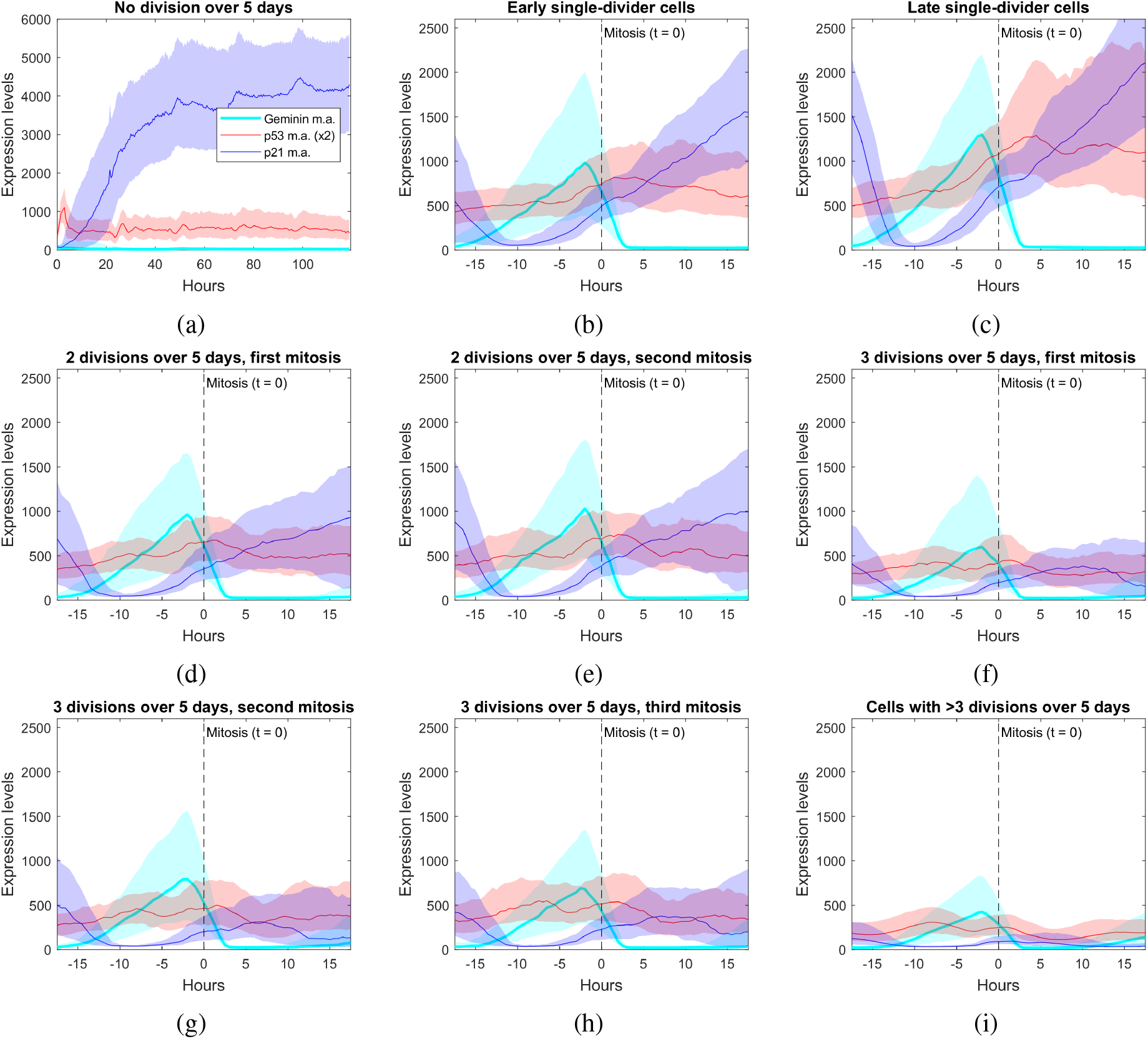
The moving averages for the expression levels of geminin, p53, and p21 are plotted and centered around mitosis (at *t* = 0) for the x-ray irradiation dataset. The middle line represents the 50 percentile value for a given relative time, while the top and bottom of the shaded region represent the 75 percentile and 25 percentile values.

Single-divider cells, shown in Fig. 7(b)-(c), undergo the process of mitosis at significantly higher values of p21 m.a. values than cells that divide more often. These cells also seem to exhibit a substantial increase of both p21 and p53 m.a. values before mitosis. These elevated levels of p53 subsist long after these cells undergo their single mitosis event. Of note, the behavior between “early single-divider cells,” defined as cells which undergo mitosis less than 3 days after exposure, exhibit distinct behavior from “late single-divider cells” (also called “escapers” in [23]). Early-single divider cells have noticeably lower levels of p21 and p53. There is also a significant jump in p53 levels in late single-divider cells near the mitosis event and afterward.

Looking in detail at cells that divided exactly twice (Fig. 7(d)-(e)) or 3 times (Fig. 7(f)-(h)), the profiles are almost identical for the same number of divisions across all three markers. There does not appear to be a significant difference in the long-term trends of p21 levels before the mitosis process has taken place and afterward, once the values are stabilized. Cells that divide more than 3 times over five days in Fig. 7(i) exhibit in aggregate the lowest levels of p21, p53, and geminin. Note that this subplot shows the behavior across all observed mitosis events, unlike the other subplots.

Cells that do not divide despite high levels of geminin are shown in Fig. 8, split into 6 clusters using k-means with correlation as the distance. These subfigures show various instances in which cells that are in the process of dividing decide to forego division or have yet to divide. These cells had the characteristics of expressing geminin levels higher than 500 a.u. for a period of more than 12.5 hours, but for which no division event was recorded. There were 175 cells in the x-ray dataset. It is relatively clear that a high level of p21 is a necessary condition at the moment that the cell reverses this decision and the levels of geminin start dropping without mitosis occurring. All these cells also have an early spike in p53, which may be due to the radiation exposure at *t* = 0. When compared to cells that do divide in the previous panel (Fig. 7), it takes longer for geminin to reach a peak, *>* 20 hrs, from the moment geminin levels start noticeably rising as opposed to 15 − 20 hrs for dividing cells. At the end of this reversion, cells return to having high m.a. values of p53 and p21, similarly to non-dividing cells as shown in Fig. 7(a).

**Figure 8:**
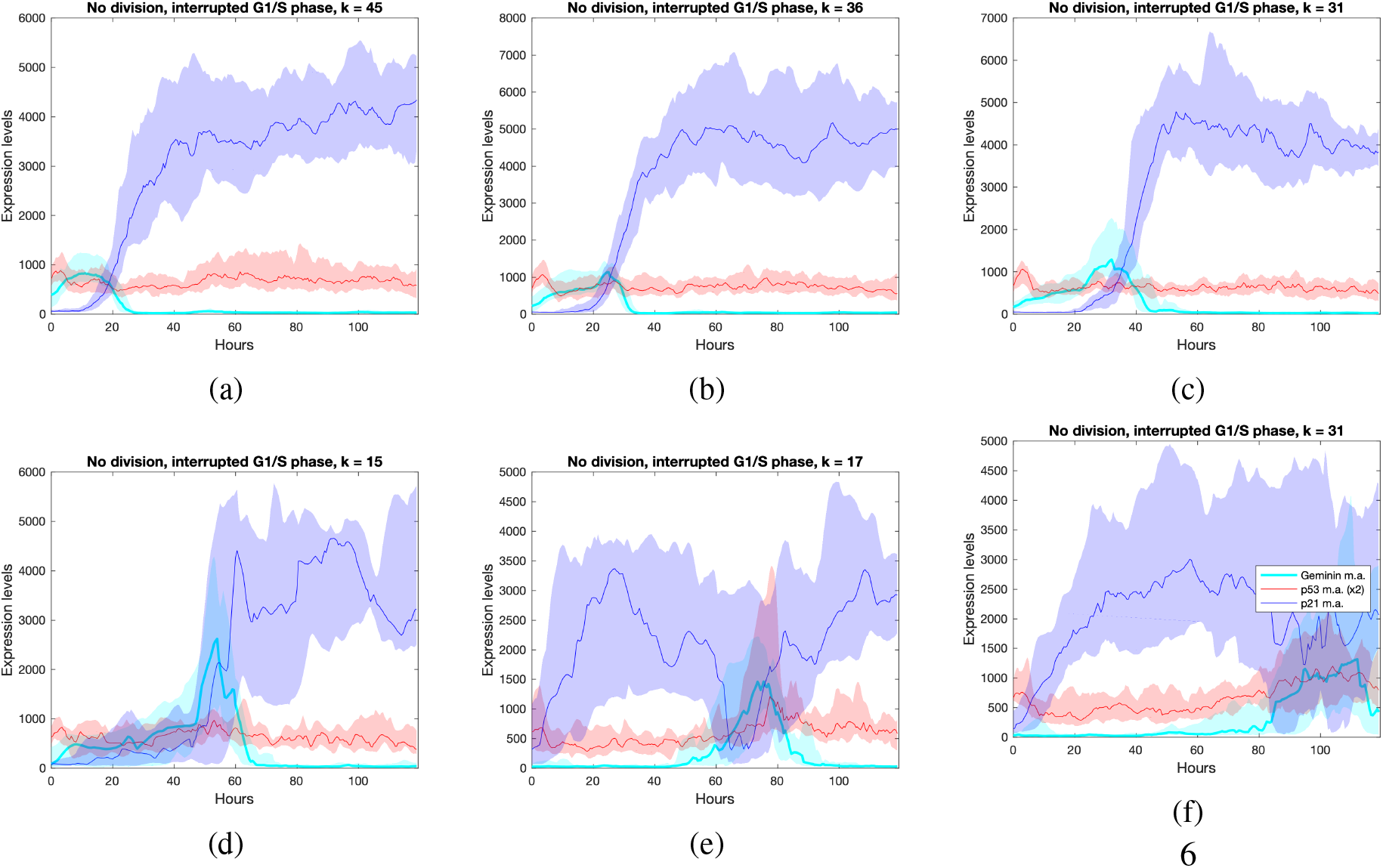
The moving averages for the expression levels of geminin, p53, and p21 are plotted and centered around mitosis (at *t* = 0) for the x-ray irradiation dataset for cells that showed signs of division, but reversed course. These cells exhibited high levels of geminin for a period of more than 12.5 hours (>500 a.u.), but did not undergo mitosis. These cells were then split into 6 clusters using k-means clustering with the distance used being correlation. The middle line represents the 50 percentile value for a given relative time, while the top and bottom of the shaded region represent the 75 percentile and 25 percentile values.

In Fig. 9, the same results as Fig. 7 are shown for the gamma radiation dataset. The geminin levels were not measured in these single cells. The m.a. trends in p21 and p53 are nearly identical between the two datasets. A difference is that raw p21 levels in the gamma dataset are of lower values than the x-ray radiation dataset.

**Figure 9:**
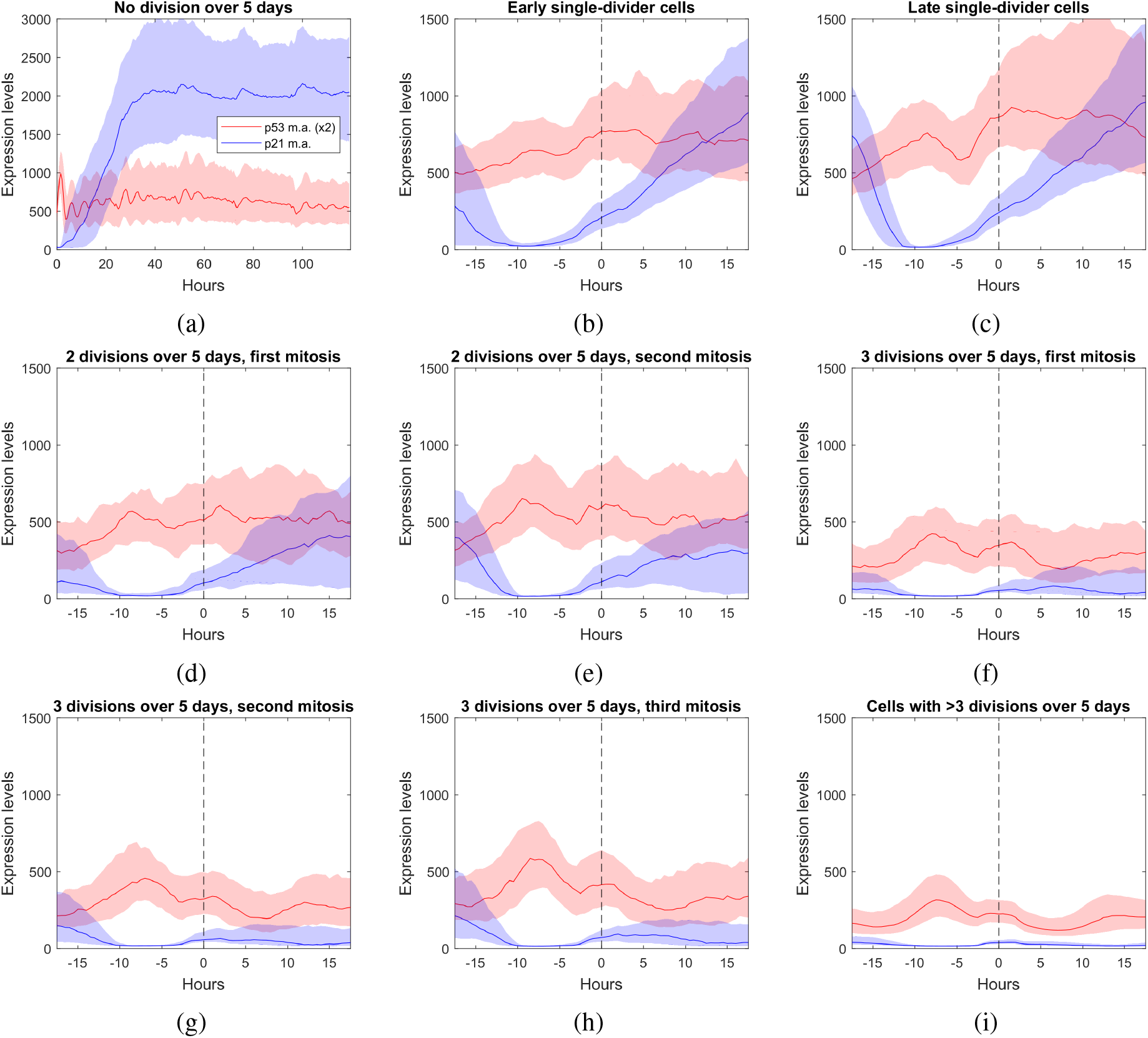
The moving averages for the expression levels of geminin, p53, and p21 are plotted and centered around mitosis (at *t* = 0) for the gamma irradiation dataset. The middle line represents the 50 percentile value for a given relative time, while the top and bottom of the shaded region represent the 75 percentile and 25 percentile values.

We should also note that the x-ray radiation profiles were very similar before and after mitosis. Looking at Fig. 9(d)-(e), one can notice elevated levels in the case of 2 divisions, after the first mitosis and before the second mitosis.

Due to p21 levels being suppressed 15-20 hours before mitosis, Fig. 10 focuses on the mean levels of p53 and p21 across all five days, but excluding the regions in which cells are actively dividing. These excluded regions were taken as the 15 hours or 30 data points leading up to a mitosis event. Looking at the distributions for mean p53 of the non-dividing regions (n.d.) in both Fig. 10(a) and Fig. 10(b), there is a correspondence between higher levels of p53 and fewer divisions. However, the difference between the mean p53 n.d. distributions of 0, 1, and 2 divisions are almost indistinguishable. Yet, it is clear that cells that do not divide are mostly treated at the most extreme radiation levels of 10 Gy for gamma radiation and 8 Gy for x-ray radiation, but this difference in the treatments is not reflected in the mean p53 n.d. values. There is also a second cluster for *>* 3 divisions over 5 days with lower mean p53 n.d. values.

**Figure 10:**
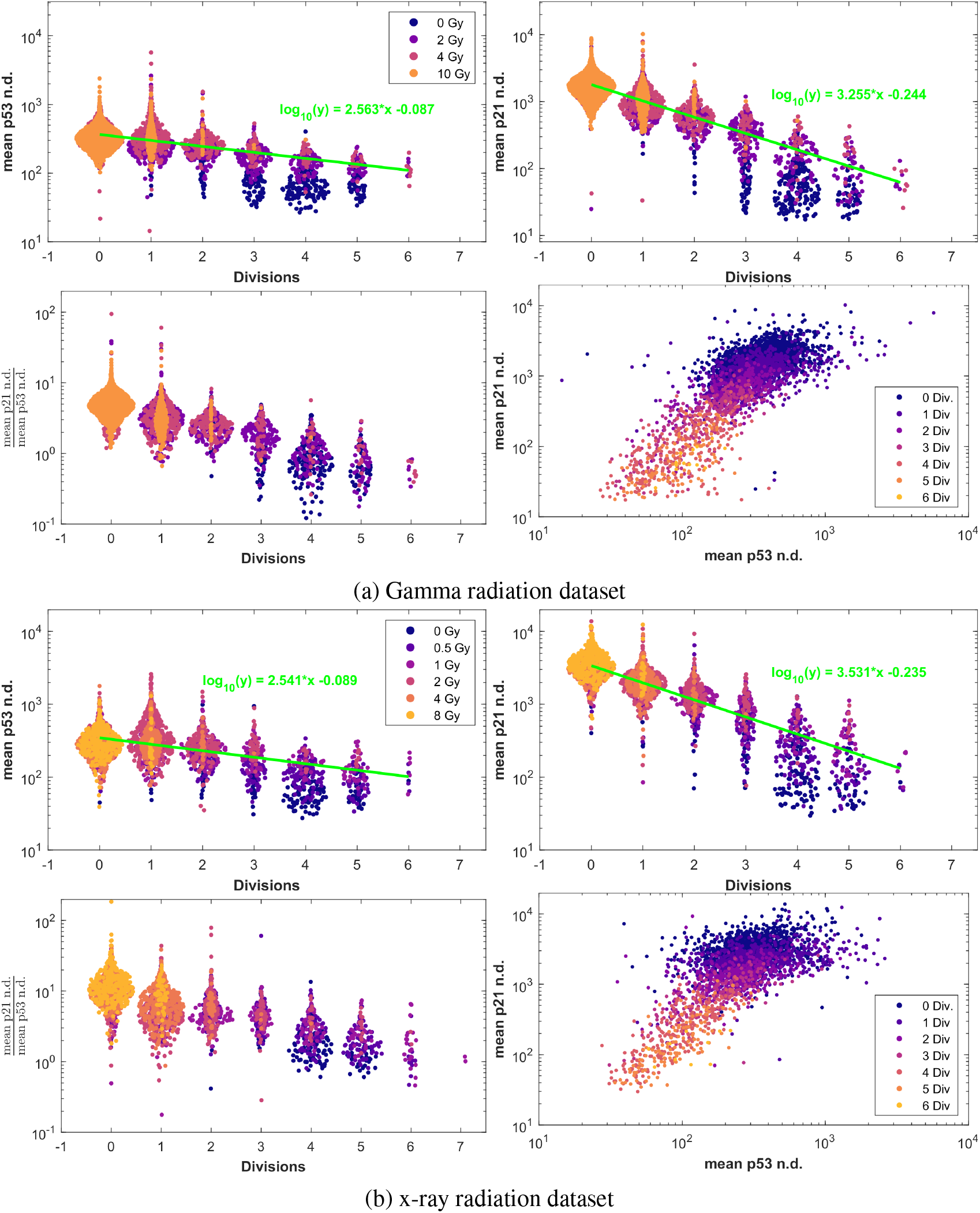
Swarm plots of the mean values of p21 and p53 for both datasets excluding the non-dividing regions where p21 levels are suppressed. The excluded regions are the 15 hours leading to a mitosis event. Each panel is broken down into mean p53 n.d. vs. number of divisions over 5 days, mean p21 n.d. vs. number of divisions, the ratio between mean p21 n.d. and mean p53 n.d. vs. number of divisions, and mean p21 n.d. vs. mean p53 n.d.

Taking a closer look at the mean p21 n.d. values in Fig. 10, there is a noticeable difference in distribution between cells that do not divide and for each additional division that these cells take over the course of the five days. This makes it clear that mean p21 n.d. levels are primarily responsible for the single cell decision making with respect to mitosis with lower values leading more divisions over the same period of time. Thus, cells do not simply undergo a binary decision of dividing or not dividing. Mean p21 n.d. levels in a given single cell strongly correlates with how frequently mitosis occurs in that cell. Plotting the mean of the non-dividing regions of p21 and p53 against one another also provides a cleaner correlation, though still noisy, between the two variables as shown in Fig. 6(a) or Fig. S1(a), where one observes higher levels of p21 in the presence of higher levels of p53.

Shown in Fig. 11 are swarm plots similar to Fig. 10, but limited to the ranges of 18-36 hours from the 5 days that the data was recorded after irradiation. The reason for picking this time frame is because it takes time for the long-term trends in p21 to settle in that may take up to a few days for some cells. Through these plots for both Fig. 11(a) and (b), we show that using these 15% of early data out of the full five days provide similar trends to those shown in Fig. 10. This provides the ability to predict the rate of division of a cell over a longer interval based on early observations because these levels of p21 tend to maintain before and after mitosis.

**Figure 11:**
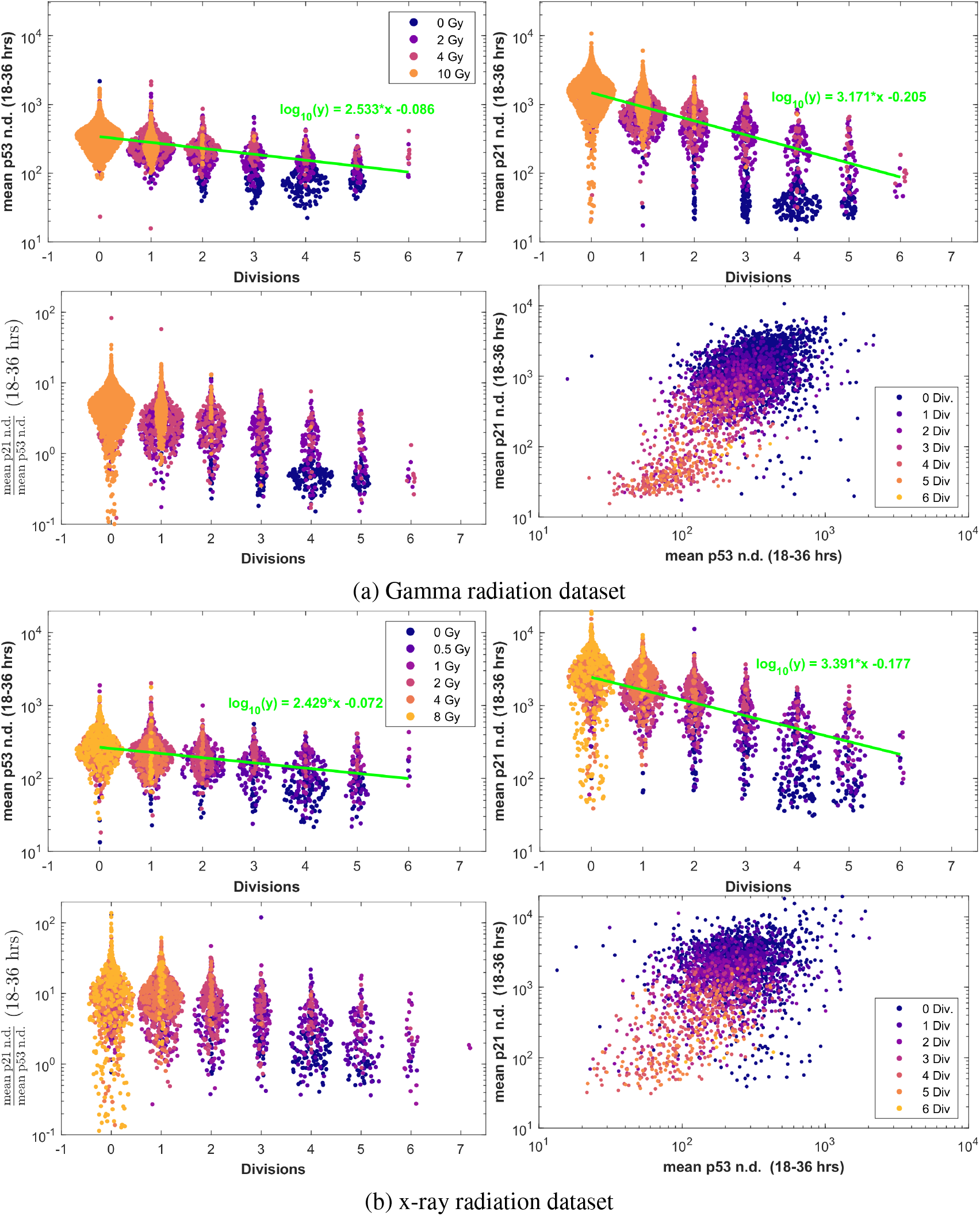
Swarm plots of the mean values of p21 and p53 for both datasets excluding the non-dividing regions where p21 levels are suppressed but limited to hours 18 to 36. The excluded regions are the 15 hours leading to a mitosis event. Each panel is broken down into mean p53 n.d. vs. number of divisions over 5 days, mean p21 n.d. vs. number of divisions, the ratio between mean p21 n.d. and mean p53 n.d. vs. number of divisions, and mean p21 n.d. vs. mean p53 n.d.

## 5 Discussion and Conclusion

Two sets of human single cells, treated with various levels of x-ray and gamma radiation, were monitored for cell division, p21, p53, and hGeminin(1-110) levels after radiation exposure [23, 29]. From the data on mitosis occurrences, it is clear that the behavior of cells within a given treatment regimen, in terms of radiation levels and type, can be quite heterogeneous with cells dividing at different rates. Using t-SNE classification with the 1-Wasserstein distance, it was found that cells can be readily clustered using the number of cell divisions they undergo over five days, using the “histograms” of p21, and to a lesser extent, p53. This indicates that p21 dynamics plays a key role in the cell decision to undergo mitosis or not, and also affects the rate at which this occurs.

While it is known that biologically, p21 is located downstream of the p53 signaling pathway, this work shows how p53 signaling is an integral part of the p21 response by analyzing the oscillatory components of the two signals. Through detrending and normalizing the two signals using the DAPS technique, it was shown how it is expected for p53 to lead p21 by about 3.5 hours and that the period of the oscillations is about 6 hours. This analysis was made possible because of the rare occurrence of mitosis in heavily damaged cells, as p53 signaling is heavily suppressed in the G1/S phases.

From studying time-lapses of the moving averages of p21 and p53, it was found that p53 trends tend to settle within hours of cell damage, while p21 values may take up to a few days to fully settle into their long-term trends. Most cells that did not divide follow a pattern of expressing very high levels of p21 that builds up from the onset of the radiation damage. However, in rarer cases, cells may be attempting to divide through the high expression of geminin and the degradation of p21, but ultimately reversed course and these cells then express very high values of p21. It is also made clear through this work that the cell decision to divide is more complex than simply the idea that cells resume division after having repaired the radiation damage. In the studied datasets, cells that had high levels of p21 before mitosis oftentimes recovered similarly high p21 levels after mitosis. Finally, we found, looking at the mean values of p21 that exclude the parts where cells are actively dividing, that the pace at which cells divide is regulated by the long-term trends of p21. This would also suggest that cells do not necessarily decide to cease division when considered too damaged, but that very high levels of p21 make it unlikely for cells to divide within the observation period of five days.

## Acknowledgments

The authors thank Professor Galit Lahav from Havard Medical School for making the data available to use in this study. This study was supported by AFOSR grant (FA9550-17-1-0435), a grant from National Institutes of Health (R01-AG048769), MSK Cancer Center Support Grant/Core Grant (P30 CA008748), and a grant from Breast Cancer Research Foundation (BCRF-17-193). CM is supported by NSF DMS grant 2111322. J.R. received support from CONACyT/Fundacion Mexico en Harvard (404476), and Harvard Graduate Merit Fellowship.

## Supplementary Materials

**Figure S1:**
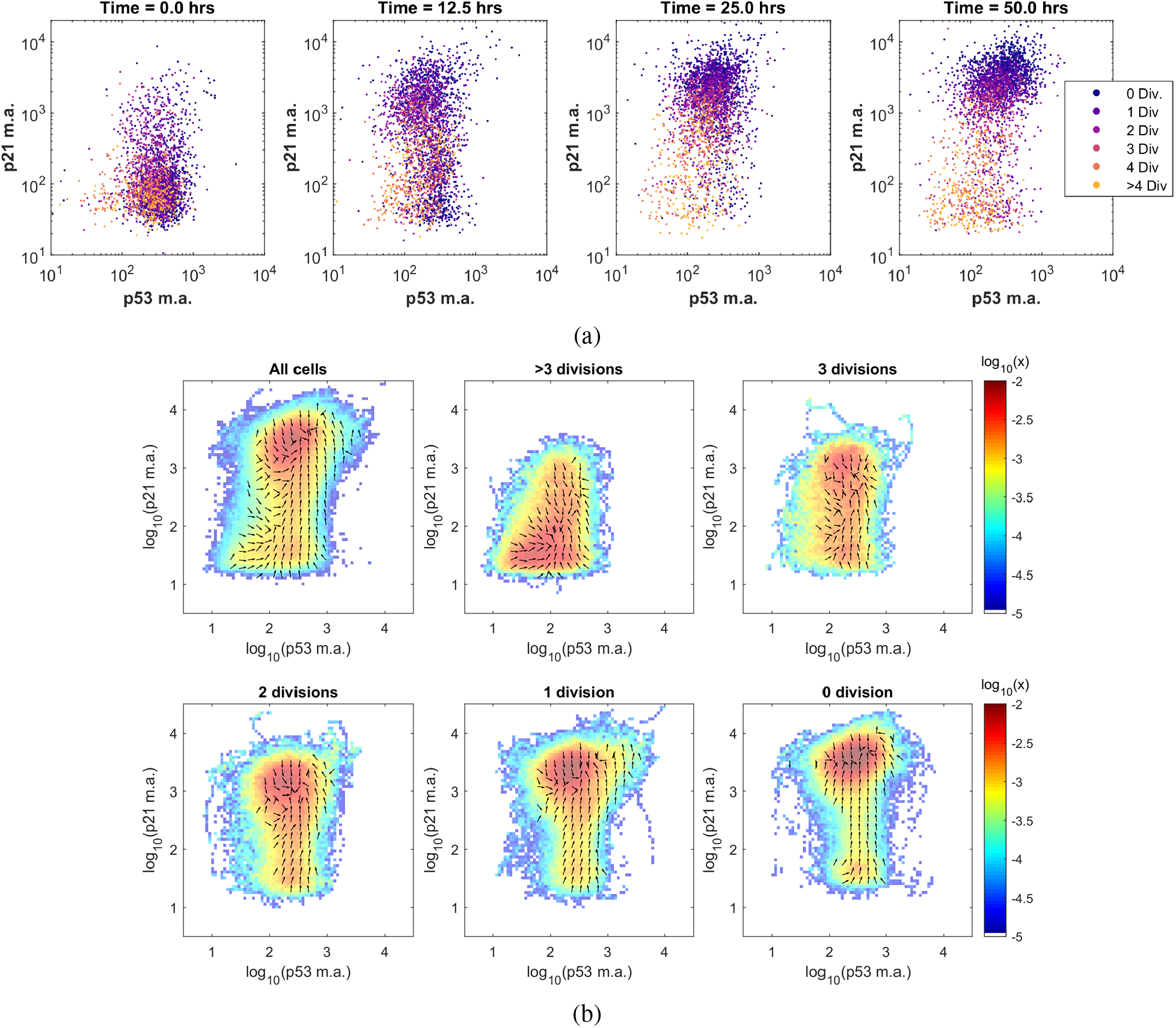
2D representations of the p21 and p53 temporal data for the x-ray radiation dataset. (a) Time lapses of the single cells of the moving average values of p21 vs. p53 at different time points of 0, 12.5, 25, and 50 hours after irradiation. The panels are broken down either by number of total divisions over five days or by irradiation conditions. (b) Occupancy density of the cells with a breakdown by the total number of undergone divisions over five days. The numerical value shown in color indicates the probability for a cell to occupy a given square of 0.1 by 0.1 in the log_10_(p21 m.a.) vs. log_10_(p53 m.a.) space in a given hour.

**Figure S2:**
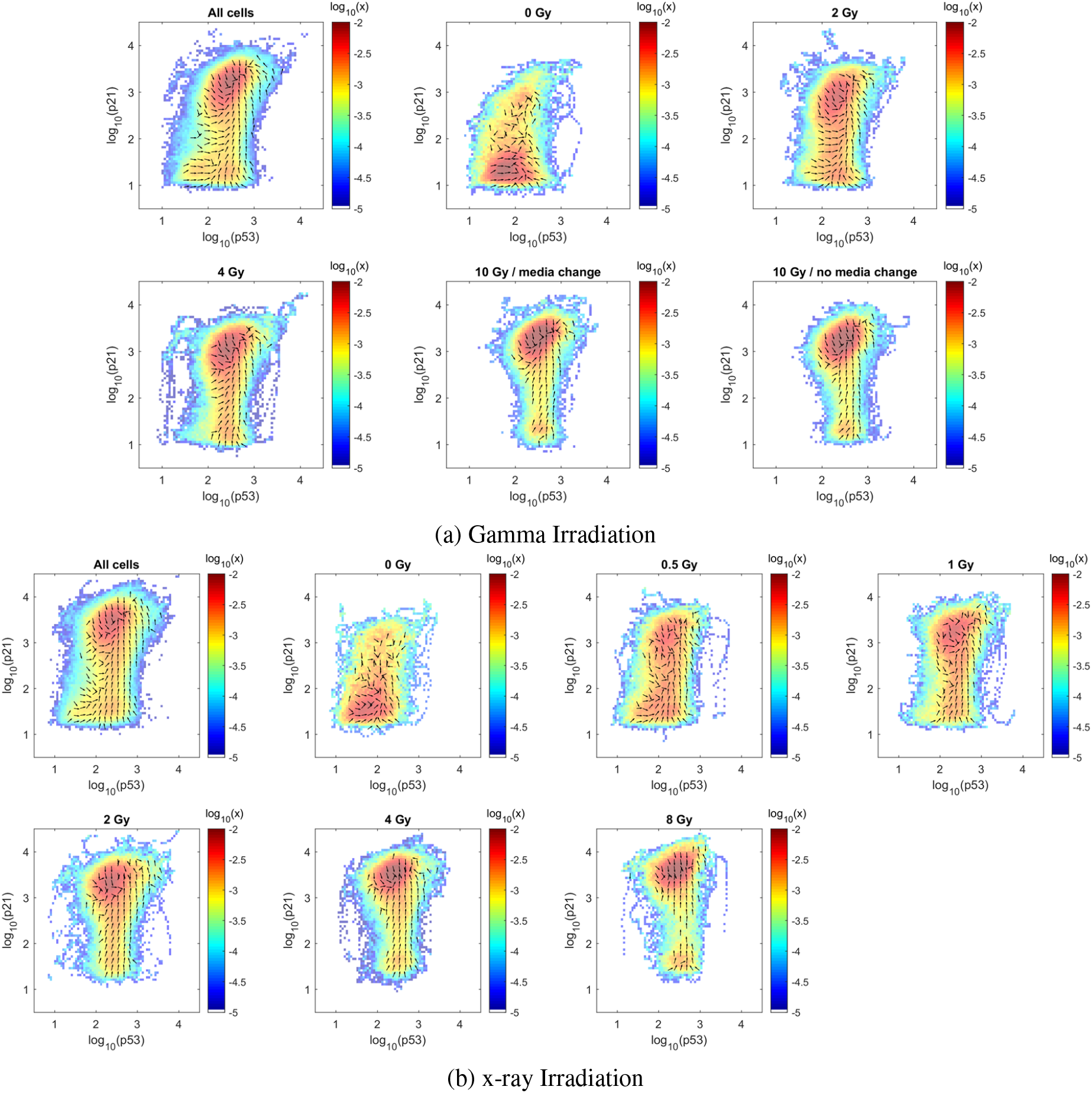
Occupancy density of the cells with a breakdown by treatment conditions for (a) the gamma- irradiation and (b) the x-ray irradiation datasets. The numerical value shown in color indicates the probability for a cell to occupy a given square of 0.1 by 0.1 in the *log*_10_(p21 m.a.) vs. *log*_10_(p53 m.a.) space in a given hour.

**Figure S3:**
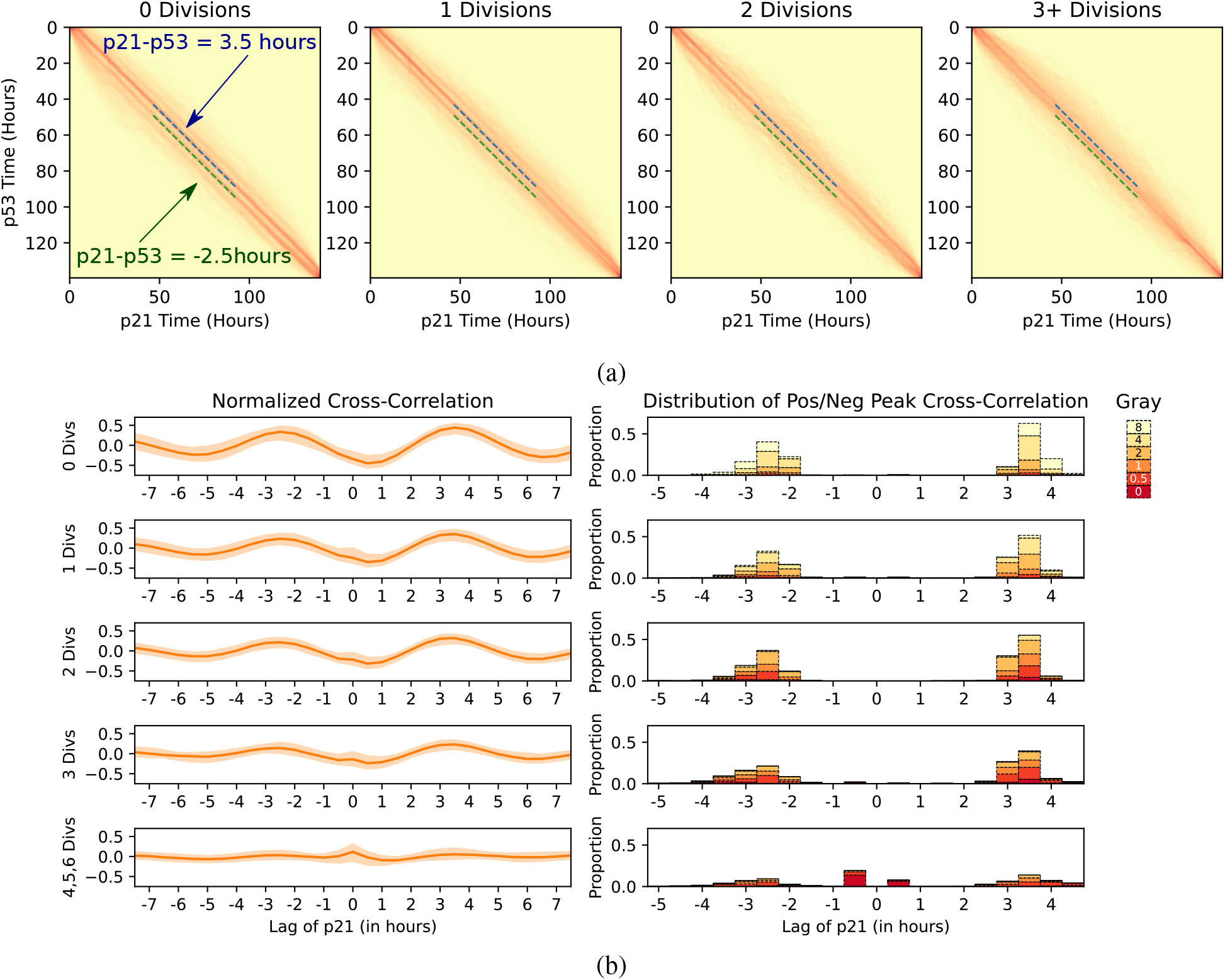
Delay results for the x-ray irradiation dataset.

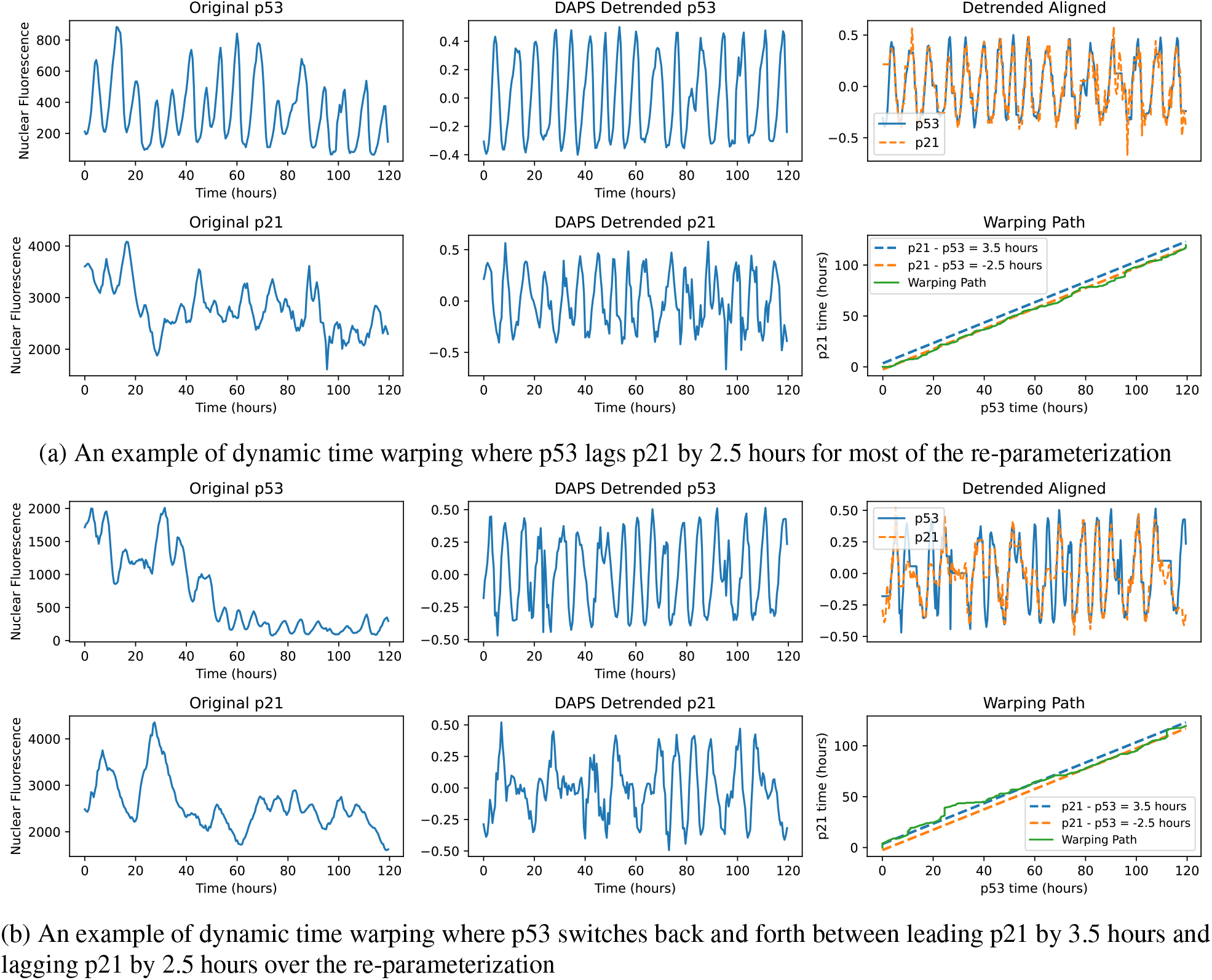

## References

[1] Ruth Lev Bar-Or, Ruth Maya, Lee A Segel, Uri Alon, Arnold J Levine, and Moshe Oren. Generation of oscillations by the p53-mdm2 feedback loop: a theoretical and experimental study. Proceedings of the National Academy of Sciences, 97(21):11250–11255, 2000.

[2] Alexis R Barr, Samuel Cooper, Frank S Heldt, Francesca Butera, Henriette Stoy, Jörg Mansfeld, Béla Novák, and Chris Bakal. Dna damage during s-phase mediates the proliferation-quiescence decision in the subsequent g1 via p21 expression. Nature communications, 8(1):1–17, 2017.

[3] Eric Batchelor, Caroline S Mock, Irun Bhan, Alexander Loewer, and Galit Lahav. Recurrent initiation: a mechanism for triggering p53 pulses in response to dna damage. Molecular cell, 30(3):277–289, 2008.

[4] Christian M Beauséjour, Ana Krtolica, Francesco Galimi, Masashi Narita, Scott W Lowe, Paul Yaswen, and Judith Campisi. Reversal of human cellular senescence: roles of the p53 and p16 pathways. The EMBO journal, 22(16):4212–4222, 2003.

[5] Donald J Berndt and James Clifford. Using dynamic time warping to find patterns in time series. In KDD workshop, volume 10, pages 359–370. Seattle, WA, USA:, 1994.

[6] Gil Bornstein, Joanna Bloom, Danielle Sitry-Shevah, Keiko Nakayama, Michele Pagano, and Avram Hershko. Role of the scfskp2 ubiquitin ligase in the degradation of p21cip1 in s phase. Journal of Biological Chemistry, 278(28):25752–25757, 2003.

[7] James Brugarolas, Chitra Chandrasekaran, Jeffrey I Gordon, David Beach, Tyler Jacks, and Gregory J Hannon. Radiation-induced cell cycle arrest compromised by p21 deficiency. Nature, 377(6549):552–557, 1995.

[8] Carl Chiarella, Xue-Zhong He, and Cars Hommes. A dynamic analysis of moving average rules. Journal of Economic Dynamics and Control, 30(9-10):1729–1753, 2006.

[9] MA Christophorou, I Ringshausen, AJ Finch, L Brown Swigart, and GI Evan. The pathological response to dna damage does not contribute to p53-mediated tumour suppression. Nature, 443(7108):214–217, 2006.

[10] Kate E Coleman, Gavin D Grant, Rachel A Haggerty, Kristen Brantley, Etsuko Shibata, Benjamin D Workman, Anindya Dutta, Dileep Varma, Jeremy E Purvis, and Jeanette Gowen Cook. Sequential replication-coupled destruction at g1/s ensures genome stability. Genes & development, 29(16):1734–1746, 2015.

[11] Manuel Collado and Manuel Serrano. Senescence in tumours: evidence from mice and humans. Nature Reviews Cancer, 10(1):51–57, 2010.

[12] Annette MG Dirac and René Bernards. Reversal of senescence in mouse fibroblasts through lentiviral suppression of p53* 210. Journal of Biological Chemistry, 278(14):11731–11734, 2003.

[13] Alejo Efeyan and Manuel Serrano. p53: guardian of the genome and policeman of the oncogenes. Cell cycle, 6(9):1006–1010, 2007.

[14] Naama Geva-Zatorsky, Nitzan Rosenfeld, Shalev Itzkovitz, Ron Milo, Alex Sigal, Erez Dekel, Talia Yarnitzky, Yuvalal Liron, Paz Polak, Galit Lahav, et al. Oscillations and variability in the p53 system. Molecular systems biology, 2(1):2006–0033, 2006.

[15] J Wade Harper, Guy R Adami, Nan Wei, Khandan Keyomarsi, and Stephen J Elledge. The p21 cdk-interacting protein cip1 is a potent inhibitor of g1 cyclin-dependent kinases. Cell, 75(4):805–816, 1993.

[16] Ygal Haupt, Ruth Maya, Anat Kazaz, and Moshe Oren. Mdm2 promotes the rapid degradation of p53. Nature, 387(6630):296–299, 1997.

[17] Do-Hyun Kim, Kyoohyoung Rho, and Sunghoon Kim. A theoretical model for p53 dynamics: identifying optimal therapeutic strategy for its activation and stabilization. Cell Cycle, 8(22):3707–3716, 2009.

[18] David P Lane. p53, guardian of the genome. Nature, 358(6381):15–16, 1992.

[19] Caroline Moosmüller, Christopher Tralie, Mahdi Kooshkbaghi, Zehor Belkhatir, Maryam Pouryahya, Jose Reyes, Joseph Deasy, Allen Tannenbaum, and Ioannis G. Kevrekidis. Periodicity scoring of time series encodes dynamical behavior of the tumor suppressor p53. IFAC-PapersOnLine, 54(9):488–495, 2021. 24th International Symposium on Mathematical Theory of Networks and Systems MTNS 2020.

[20] Caterina Nardella, John G Clohessy, Andrea Alimonti, and Pier Paolo Pandolfi. Pro-senescence therapy for cancer treatment. Nature Reviews Cancer, 11(7):503–511, 2011.

[21] K Wesley Overton, Sabrina L Spencer, William L Noderer, Tobias Meyer, and Clifford L Wang. Basal p21 controls population heterogeneity in cycling and quiescent cell cycle states. Proceedings of the National Academy of Sciences, 111(41):E4386–E4393, 2014.

[22] Svetlozar T. Rachev and Ludger Rüschendorf. Mass Transportation Problems: Volume I: Theory. Probability and its Applications. Springer, Berlin, 1998.

[23] José Reyes, Jia-Yun Chen, Jacob Stewart-Ornstein, Kyle W Karhohs, Caroline S Mock, and Galit Lahav. Fluctuations in p53 signaling allow escape from cell-cycle arrest. Molecular cell, 71(4):581–591, 2018.

[24] Asako Sakaue-Sawano, Hiroshi Kurokawa, Toshifumi Morimura, Aki Hanyu, Hiroshi Hama, Hatsuki Osawa, Saori Kashiwagi, Kiyoko Fukami, Takaki Miyata, Hiroyuki Miyoshi, et al. Visualizing spatiotemporal dynamics of multicellular cell-cycle progression. Cell, 132(3):487–498, 2008.

[25] Hiroaki Sakoe and Seibi Chiba. Dynamic programming algorithm optimization for spoken word recognition. IEEE transactions on acoustics, speech, and signal processing, 26(1):43–49, 1978.

[26] Sabrina L Spencer, Steven D Cappell, Feng-Chiao Tsai, K Wesley Overton, Clifford L Wang, and Tobias Meyer. The proliferation-quiescence decision is controlled by a bifurcation in cdk2 activity at mitotic exit. Cell, 155(2):369–383, 2013.

[27] Jacob Stewart-Ornstein and Galit Lahav. p53 dynamics in response to dna damage vary across cell lines and are shaped by efficiency of dna repair and activity of the kinase atm. Science signaling, 10(476), 2017.

[28] Sylvanie Surget, Marie P Khoury, and Jean-Christophe Bourdon. Uncovering the role of p53 splice variants in human malignancy: a clinical perspective. OncoTargets and therapy, 7:57, 2014.

[29] Michael Tsabar, Caroline S Mock, Veena Venkatachalam, Jose Reyes, Kyle W Karhohs, Trudy G Oliver, Aviv Regev, Ashwini Jambhekar, and Galit Lahav. A switch in p53 dynamics marks cells that escape from dsb-induced cell cycle arrest. Cell reports, 32(5):107995, 2020.

[30] Laurens Van der Maaten and Geoffrey Hinton. Visualizing data using t-sne. Journal of machine learning research, 9(11), 2008.

[31] Alexei Vazquez, Elisabeth E Bond, Arnold J Levine, and Gareth L Bond. The genetics of the p53 pathway, apoptosis and cancer therapy. Nature reviews Drug discovery, 7(12):979–987, 2008.

[32] Cédric Villani. Topics in Optimal Transportation. American Mathematical Soc., 2003.

[33] Cédric Villani. Optimal Transport: Old and New, volume 338. Springer Science & Business Media, 2008.

[34] Bert Vogelstein, David Lane, and Arnold J Levine. Surfing the p53 network. Nature, 408(6810):307–310, 2000.

[35] J Wagner, L Ma, JJ Rice, W Hu, AJ Levine, and GA Stolovitzky. p53-mdm2 loop controlled by a balance of its feedback strength and effective dampening using atm and delayed feedback. IEE Proceedings-Systems Biology, 2(3):109–118, 2005.

[36] Yue Xiong, Gregory J Hannon, Hui Zhang, David Casso, Ryuji Kobayashi, and David Beach. p21 is a universal inhibitor of cyclin kinases. Nature, 366(6456):701–704, 1993.

[37] Wen Xue, Lars Zender, Cornelius Miething, Ross A Dickins, Eva Hernando, Valery Krizhanovsky, Carlos Cordon-Cardo, and Scott W Lowe. Senescence and tumour clearance is triggered by p53 restoration in murine liver carcinomas. Nature, 445(7128):656–660, 2007.

[38] Tongli Zhang, Paul Brazhnik, and John J Tyson. Exploring mechanisms of the dna-damage response: p53 pulses and their possible relevance to apoptosis. Cell cycle, 6(1):85–94, 2007.

